# Schrödinger’s Cat in Simulations in Genome-wide Association Studies

**DOI:** 10.1101/2023.03.22.533184

**Authors:** Mustafa Ismail Ozkaraca, Albert Tenesa

## Abstract

Simulations are essential components for testing methods or assumptions in genome-wide association studies (GWAS). We show through theory and examples that markers can be both causal and non-causal simultaneously, resulting in a paradox, had simulation not being run properly. We developed a software (ParaPheSim – a paradoxical phenotype simulator) that allows users to create different traits whose genetic correlation is exactly 1 and yet they have no genetic overlap.

---

Genome-wide association studies (GWAS) have identified associations between genotypes and phenotypes for large number of complex traits and diseases over the past decade or so [1, 2]. One indispensable component of this genomic data science era is simulations. Whether it is being used for benchmarking purposes of a new statistical software [3–7] or setting up an experimental design [8] or else, statistical inference procedures based on computer simulations can be very useful for we know all underlying parameters to be estimated, a scenario which is impossible in studies based on real data. However, unfortunately, simulations can be misinterpreted if not being applied appropriately.

One subtle assumption about simulations is that generated data have to be uniquely identified. For example, if we are to simulate a phenotype from a given genotype information, we need to make sure that there is a unique set of causal markers, chosen from genotype data, that underpins the genetic component of simulated phenotype uniquely. Since otherwise different set of causal markers can generate the same phenotype and we arrive a paradox of defining a variant to be causal and non-causal simultaneously (Figure 1).

**Figure 1.**
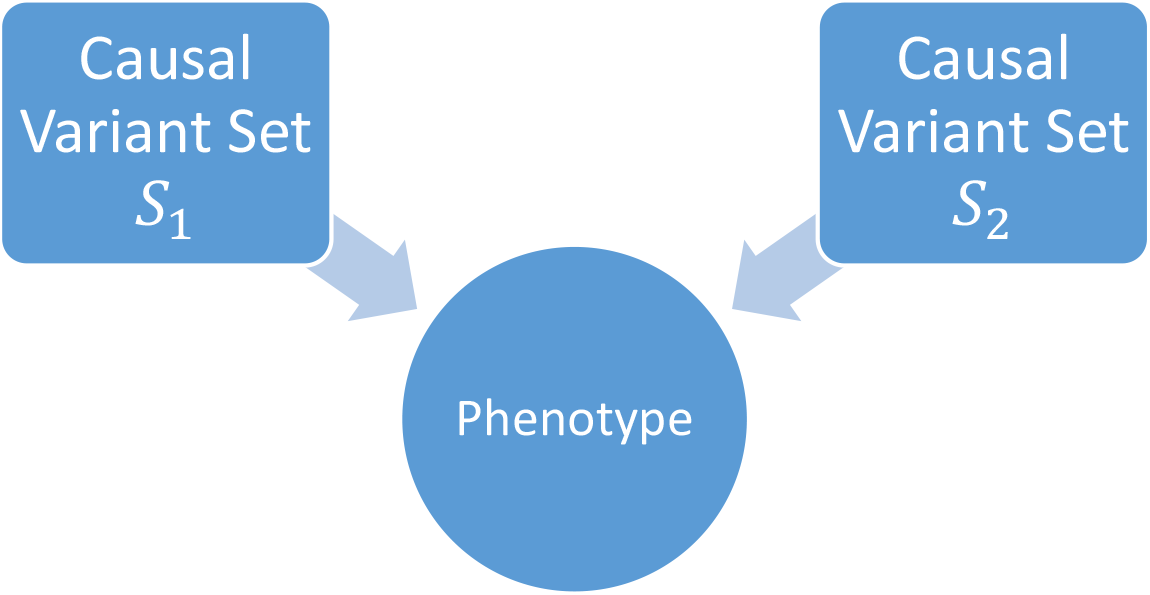
In the case of different set of causal variants generates same phenotype, a variant in *S*_1_\*S*_2_ is causal (because it is in *S*_1_ and non-causal (because it is not in *S*_2_).

To explain the enigma further with a toy data set, consider the genotype matrix G given below. For brevity, we consider an unstandardized matrix but the same logic holds even if *G* is replaced with a standardised matrix. It contains information about 4 variants (columns) for 4 individuals (rows).

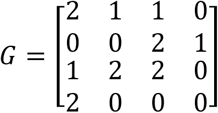

Suppose two independent groups want to generate a continuous trait from *G.* Group-1 assigns effect sizes *w*_1_ = [0 0.25 0 0.5] for variants (i.e. Variant-2 and Variant-4 are only causals with effect sizes 0.25 and 0.5 respectively) and group-2 assigns *w*_2_ = [0 0 0.25 0]. Assume further they both generate phenotypes by the formulas

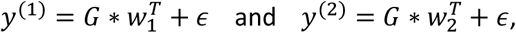

where *ϵ* denote (same) random noise and *y*^(*i*)^ denote simulated phenotype by group *i*. If, for simplicity, we ignore the random noise terms, then we get

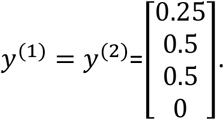

That is, both groups have generated the same phenotype by chance. These phenotypes, whenever interpreted as different traits, have genetic correlation 1, though they have no genetic overlap. Finally, suppose both groups implement the same GWAS model and arrive to the same summary results, where both groups find that Variant-2 and Variant-4 are significant and the rest are not significant, under some threshold *α*. While researchers in group-1 interpret this conclusion as their model performing well and produced no false associations, group-2 interprets it differently that the model performed poorly and produced two false positives (Figure 2). While both groups have used the same inputs, they have arrived different conclusions. This is a paradox, due to the discrepancies that Variant- 3 is causal from Group-2 point of view but is non-causal from Group-1 point of view, a dilemma reminding Schrödinger’s experiment [9].

**Figure 2.**
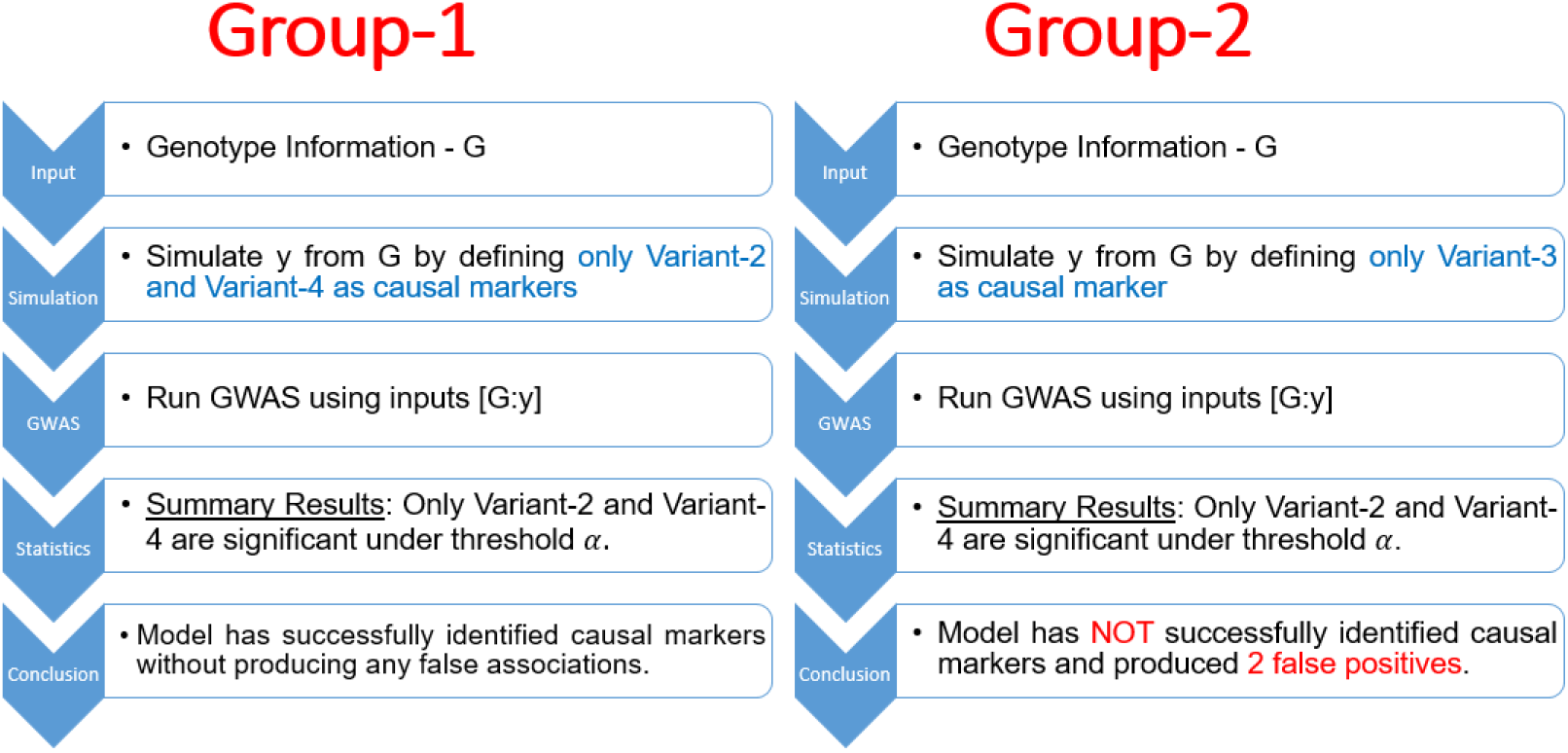
Two independent research groups generate same phenotype even though they selected different variants to be causal. They run the same GWAS independently, and Group-1 claims the model performing well whereas Group-2 concludes it performs poorly.

The fundamental source of this dilemma is that Variant-2, Variant-3 and Variant-4 form a linearly dependent set. In particular,

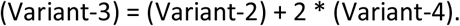

Unfortunately, the outlined issue is guaranteed to occur whenever N (number of individuals) is less than M (number of variants to be tested) (**Online Methods**). In fact, we have proved that this issue occurs at least 2^*M*–*N*^ different times in every single GWAS simulation whenever *N* < *M* (**Online Methods**).

We refer to this problem as the *input-dimension paradox.* Performance for classification of multiple hypotheses tests (true/false positives/negatives) can be vague in existence of the paradox due to same markers being both causal and non-causal simultaneously. We have created a software to show how vital this paradox can be. Given a user-choice of set of variants to be defined as causal markers, our tool simulates a continuous phenotype *y* under the model *y* = *g* + *e*, and then finds a number (given by user input) of different sets of mutually disjoint variants that can be replaced as causal markers (with different associated effect sizes) to generate the same component g. Our tool is publicly available, and includes an example of a continuous trait with 21 different representations of its genetic component.

For dichotomous traits, the anomaly is double worse because a disease simulated from a genotype does not hold information about which diseased individual has higher (or lower) underlying genetic liability. For instance, suppose we simulate a disease from the continuous trait generated above such that a person is labelled as diseased if its (continuous) phenotype value is > 0.4. This would induce a binary trait 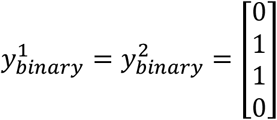. Suppose further that some other research group, group-3, wants to simulate a continuous trait using *G* by assigning effect sizes [0.05 0 0.25 0], and using the same formula as the other two groups with ignoring the random noise (for brevity). Then they generate a continuous trait 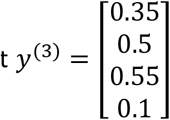 which would induce the same binary trait above, whenever one uses the same threshold > 0.4 in order to define diseased individuals. As such, by labelling even an additional variant as causal on underlying liability model (Variant-1), one can still be able to generate the same binary trait. This causes even more critical disputes about which set of variants is causing an individual to be diseased.

In order to define a unique set of causal markers for a continuous simulated trait, according to linear algebra theory, we need to have both:

A. No causal marker can be written as a linear combination of other causal variants.
B. No causal marker can be written as a linear combination of non-causal variants. (**Online Methods**)

Satisfying both conditions assures that the underlying genetic component of the simulated trait is defined uniquely and causal markers are the same for every analysis and for each group that uses the same datasets (phenotype and genotype). However, as in almost all statistical analyses, simulating one single phenotype based on a user’s choice set of (causal) variants is not recommend, because the random noise (or environmental factors) can be rearranged to cause the same paradox outlined above (i.e. a different set of variants can still be defined to be causals by rearranging the error term). For that, simulating multiple traits (we suggest at least 20 traits) with the exact same genetic component is required.

In case of binary traits, the underlying causal markers cannot be guaranteed to be defined uniquely because we are discretising individuals (i.e. labelling them by only 0 or 1) during the simulation process and ignoring the information that some people have higher disease liability than others.

Condition (A) indicates that we must have *M* ≤ *N* and choose a small subset of variants to test whenever *N* << *M*, and condition (B) implies that non-causal variants have to be linearly independent from causal markers. In practice, these conditions are often not preferred as researchers wish to test associations for a large(r) number of variants. Therefore, for practical purposes, we suggest to shift the main focus on such simulations to other measures based on effect size estimates rather than detecting associations, which leads to vague conclusions under the presence of input-dimension paradox, such as evaluating performance of predicting phenotypes under cross-validation scheme by estimating effect sizes on train data and predicting phenotypes on test data using trained parameters.

## Supporting information

All data and code used in the manuscript

## Online Methods

### 1. Fundamental Source of the Paradox

Recall that for a given matrix *A* of dimension *n* × *m, rank* (*A*) ≤ *min* {*n, m*}.

Suppose *G* is a genotype matrix (either standardised or non-standardised) of dimensions *n* × *m* so that *rank* (*G*) = *r* and *n* < *m*. There exists a matrix *S* whose columns are chosen from column vectors of *G* and has dimension *n* × *r* so that *rank* (*S*) = *rank* (*G*) = *r*. Assume that *C* denote a matrix with dimension *n* × *k*, whose columns corresponds to variants and are chosen (by a user) from column vectors of *G* to be causal markers for a continuous trait to be simulated. In matrix form,

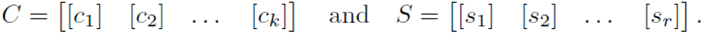

Since *rank* (*S*) = *rank* (*G*) = *r* and column vectors of both *C* and *S* are also column vectors of *G*, every column vector of *C* is a linear combination of column vectors of *S.* Precisely, we have

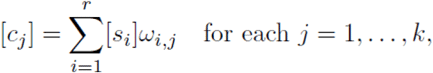

where *ω_ij_* is a scalar for each *i,j.* Equivalently,

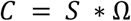

where Ω is a matrix with dimension *r* × *k* and *ω_ij_* denote its element in *i*-th row and *j*-th column, for each *i* ∈ {1,…, *r*} and *j* ∈ {1,…, *k*}. Hence, whenever a continuous trait is simulated under the simple genetic additive model *y* = *g* + *e* so that

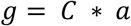

for some row vector *a* which corresponds to true effect sizes of causal markers defined by *C* (i.e. columns of *C*), a different representation of *g* can be given by

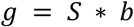

where *b* = Ω * *a* can be interpreted as associated effect sizes of variants (i.e. columns) of *S*. As such, a different matrix *S* corresponding to causal markers, other than *C,* can be defined in order to generate the very same phenotype (using the same *e* component).

#### Remark

We have not put any condition on neither *G* or other defined matrices. In our tool, we use standardised genotype matrix (similar to [10]) (See Section 4). However, the logic holds even non-standardised matrices are used.

#### Computing Ω Efficiently

We have outlined that Ω can be constructed by writing each column of *C* in terms of columns of *S.* This can be computationally cumbersome especially whenever number of variants is a large number. Alternatively, we can compute Ω as follows

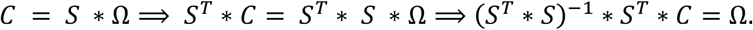

Note that whenever *S* consists of linearly independent column vectors, *S^T^* * *S* is an invertible matrix. This is computationally efficient, though inverting the matrix *S^T^* * *S* could require large amount of memory depending on the rank *r* of genotype matrix *G*.

#### Finding *S*

The fundamental ingredient in the discussion above is the matrix *S.* By selecting subsets of columns of *G,* one can always construct a matrix *S* such that *rank* (*S*) = *rank* (*G*). In theory, existence of at least one such matrix is guaranteed. Here, we outline an algorithm how to find such a matrix *S*.

**Algorithm 1.**
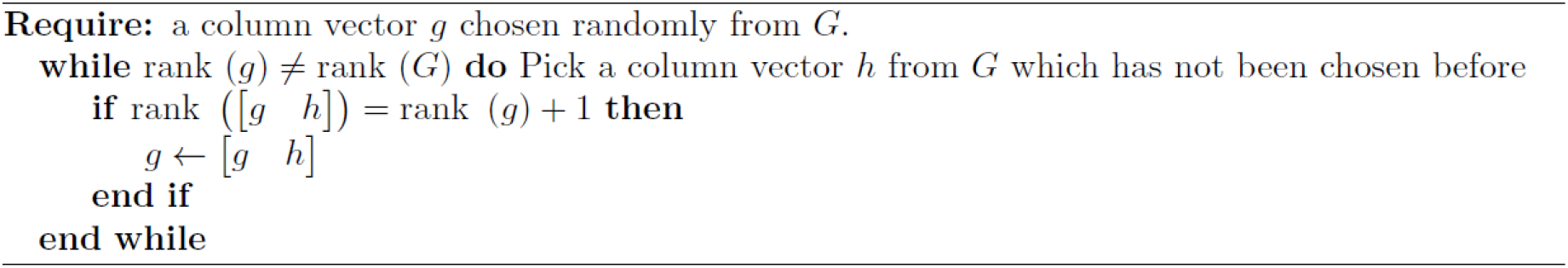
An algorithm to construct *S*

### 2. Number of Different Representations of Genetic Components

We have showed in Section 1 that for any given set of causal markers (corresponds to matrix *C*), there exists another set of variants (corresponds to matrix *S*) that can also be interpreted as the set of causal markers of the same trait. We prove next that any matrix *T* of the form *T* = [*S A*] where *A* is a matrix whose columns are chosen randomly from column vectors of *G*, can also be replaced by *S* to define even another different set of variants as the underlying causal markers.

With the same notations as before, let *T* = [*S A*] be a matrix of dimension *n* × (*r* + *a*) whose columns are column vectors of *G.* Since columns of *A* are also column vectors of *G,* we have

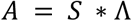

for some matrix Λ of dimension *r* × *a*. Define randomly a matrix *U* of dimension *a* × *k* such that Λ * *U* is not the zero-matrix (of dimension *r* × *k*). Such a matrix exists. Define further *V* = Ω + Λ* *U*. Then

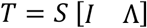

where *I* denote identity matrix of dimension *r* × *r*. This leads to,

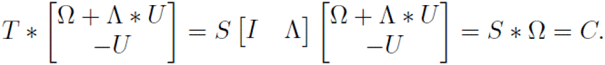

Hence,

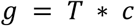

where 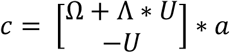.

We proved that any matrix containing *S* can also induce a (different) set of causal markers for explaining the same genetic component *g*.

Note that there are **2^*M*–*N*^ matrices** induced from columns of *G* that contains (columns of) *S.* For the case of M=1,000,000 and N=500,000, 2^*M–N*^ is a 150,515 digits and about 40 pages long number.

### 3. Constructing Unique Set of Causal Markers for the Genetic Component

We have explained in previous sections that genetic components are not uniquely defined whenever *n* < *m*. We show now mathematical requirement for how to define genetic components uniquely.

#### Theorem

Suppose *G* denote a genotype matrix of dimension *n* × *m* and [*u*_1_, [*u*_2_], …, [*u_m_*] are its column vectors of dimension *n* × 1. Suppose further we choose first *r* column vectors of *G* (for simplicity) to be causal markers to simulate a continuous phenotype under the model *y* = *g* + *e* where *g* = *C* * *a* such that *C* = [[*u*_1_], [*u*_2_],…,[*u_r_*]] and *a* is a column vector of dimension *r* × 1 which corresponds to effect sizes assigned on column vectors [*u*_1_], [*u*_2_],…,[*u_r_*]. Then there is no matrix *Y* with *Y* ≠ *C* and whose columns are chosen from column vectors of *G* such that *g = Y * b* for some row vector *b* if

i. *rank* (*C*) = min {*n, r*}, *and*
ii. *span*({[*u_i_*] : *i* ∈ {*r* + 1,…, *m*}}) ⊆ *span* ({[*u_i_*]: *i* ∈ {1,…, *r*}})^⊥^.

#### Proof

We have that *g* ∈ *span*({[*u_i_*]: *i* ∈ {1,…,*r*}}) because *g* = *C* * *a*. Therefore,

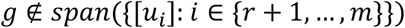

by (ii). That is, *g* can only be defined using the column vectors [*u*_1_,…,[*u_r_*]. In addition, by (i), we have that the set of column vectors of *C* is a linearly independent set and therefore every element in *span*({[*u_i_*]: *i* ∈ {1,…,*r*}}) is uniquely defined. Hence, such matrix *Y* with *Y* ≠ *C* cannot exists. QED.

### 4. ParaPheSim

For a given matrix *C* to be defined as of genotype values of causal markers and consisting of standardised vectors, we generate a continuous phenotype *y* under the simple additive model *y* = *g* + *e* where

i. *g* = *C* * [*a*] for some column vector [*a*] such that 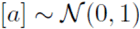, and
ii. 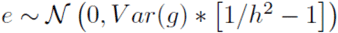 where *h*^2^ is SNP-heritability parameter given by user input.

Given a user-input integer *n*, our tool then creates *n* many different representations of the same component *g* using different set of markers. In a special case of HapMap3 dataset [11], we further force those different sets of markers to be mutually disjoint [12].

## Acknowledgements

The work was funded by Roslin Institute Strategic Programme Grants from the BBSRC (BBS/E/D/10002070 and BBS/E/D/30002275) and Health Data Research UK (references HDR-9004 and HDR-9003). For the purpose of open access, the author has applied a CC-BY public copyright licence to any Author Accepted Manuscript version arising from this submission.

