## Supplementary material for "Schrödinger’s Cat in Simulations in Genome-wide Association Studies": All data and code used in the manuscript: replicate.genetic_data.sh

```
#!/bin/bash

#################################################################
########################### VARIABLES ###########################
#################################################################

# Change your directory to a folder of your preference before implementing the following steps

dir_user=`pwd`  # current directory
git clone https://git.ecdf.ed.ac.uk/iozkarac/paraphesim
## Following are Sample and Variant IDs to generate the same dataset
shuffled_snps=${dir_user}/paraphesim/data/shuffled.snps.100k # input path of shuffled snps -> M=100.000
shuffled_ids=${dir_user}/paraphesim/data/shuffled.ids.1k # input path of shuffled ids -> N=1.000

#################################################################
########################### MODULES #############################
#################################################################

# You need to download plink2.0. 
# Please check https://www.cog-genomics.org/plink/ for download options..

#################################################################
########## DOWNLOAD AND EXTRACT THE DATA TO A FOLDER ############
#################################################################

cd ${dir_user}
wget https://ftp.ncbi.nlm.nih.gov/hapmap/genotypes/hapmap3_r3/plink_format/hapmap3_r3_b36_fwd.consensus.qc.poly.map.gz
wget https://ftp.ncbi.nlm.nih.gov/hapmap/genotypes/hapmap3_r3/plink_format/hapmap3_r3_b36_fwd.consensus.qc.poly.ped.gz

gzip -d hapmap3_r3_b36_fwd.consensus.qc.poly.map.gz
gzip -d hapmap3_r3_b36_fwd.consensus.qc.poly.ped.gz

#################################################################
####################### FILTER THE DATA #########################
#################################################################

geno=${dir_user}/hapmap3_r3_b36_fwd.consensus.qc.poly
out1=${dir_user}/temp1
out2=${dir_user}/temp2
out_final=${dir_user}/hapmap3.subset

plink --file ${geno} --make-bed --out ${geno}
plink2 --bfile ${geno} --maf 0.01 --hwe 1e-15 --geno 0.01 --mind 0.01 --chr 1-2 --indep-pairwise 1000 100 0.9 --snps-only --write-samples --write-snplist --out ${out1}
plink2 --bfile ${geno} --make-bed --keep ${out1}.id --extract ${out1}.prune.in --out ${out2}
plink2 --bfile ${geno} --keep ${shuffled_ids} --extract ${shuffled_snps} --make-bed --out ${out_final}

rm ${out1}*
rm ${out2}*
```
