## Supplementary material for "Schrödinger’s Cat in Simulations in Genome-wide Association Studies": All data and code used in the manuscript: paraphesim.py

```
"""
ParaPheSim
"""

# Modules

import argparse
import os
import sys
import time
import datetime

import pandas as pd
import numpy as np
import random
from pysnptools.snpreader import Bed


# Argparse
parser = argparse.ArgumentParser(prog="ParaPheSim", description="A Paradoxical Phenotype Simulator", epilog= "For more information please see https://git.ecdf.ed.ac.uk/iozkarac/para-phe-sim")
parser.add_argument("--hapmap", help="Example Usage: --hapmap <parameters>", action='store_true')
parser.add_argument("--paradox", help="Example Usage: --paradox <parameters>", action='store_true')
parser.add_argument("--geno", help="Example Usage: --geno <genotype_file_path>", metavar="<genotype_file_path>")
parser.add_argument("--initial_causal", help="Example Usage: --initial_causal <path>. Note that format has to be as follows: no header & second column contains variants info", metavar="<files_path_prefix>")
parser.add_argument("--replicate", help="Example Usage: --replicate 2.")
parser.add_argument("--index", help="Example Usage: --index <path>")
parser.add_argument("--h_sqrt", help="Example Usage: --h_sqrt 0.3")
parser.add_argument("--out", help="output help", default="paraphesim", metavar="<output file name>")

initial_message='''**********************************
A Paradoxical Phenotype Simulator
Version: v.0.0
**********************************

This version does only one job: simulate a continuous trait, under the model y = g + e, with given parameters so that the generated phenotype y has multiple different representations for the underlying genetic component g.

Additional features might be added later..

A manual is available here:
https://git.ecdf.ed.ac.uk/iozkarac/PARAPHESIM
__________________________________
'''

if __name__ == "__main__":
	args = parser.parse_args()
	print(initial_message)  
	assert args.out is not None, "Output path is required.."

	input_vars = vars(parser.parse_args())
	with open(args.out + ".log", "w") as f:

		f.write("ParaPheSim v.0.0")
		f.write("\nAnalysis Start Time: %s" % datetime.datetime.now().strftime("%A - %d %B %Y - %H:%M:%S"))   
		start_time=time.time()
		f.write("\nWorking Directory: " + str(os.getcwd()))
		f.write("\nInput Variables:\n")
		for key in input_vars:
			if input_vars[key]!= None and type(input_vars[key])!=bool:
				f.write("\t--%s %s\n" % (str(key), str(input_vars[key])))
			elif input_vars[key]!=vars(parser.parse_args([]))[key] and type(input_vars[key])==bool:
				f.write("\t--%s \n" % str(key))

	if args.hapmap:
		print('\nSimulating a continuous trait...')

		assert args.geno is not None, "Path for bed file is required.."
		assert args.initial_causal is not None, "List of causal variants is required.."
		assert args.replicate is not None, "Replicate is the number of distinct representations of the underlying genetic component. It is required.."
		assert args.h_sqrt is not None, "Heritability parameter is required.."

		her = float(args.h_sqrt)
		rep = int(args.replicate)
		bed_path = args.geno + '.bed'
		fam_path = args.geno + '.fam'
		bim_path = args.geno + '.bim'
		causal_path = args.initial_causal
		index_path = args.index
		out_causal = args.out + ".causal_%s"
		out_pheno = args.out + ".pheno"
		out_prs = args.out + ".prs_%s"

		snpreader = Bed(bed_path, count_A1=True)

		print('Loading variants..')
		df_variants = pd.read_csv(causal_path, index_col=None, header=None, delim_whitespace=True)   # No column and Second column is the variant info
		variants = df_variants[1].tolist()
		del df_variants

		G_temp = snpreader[:,snpreader.sid_to_index(variants)]
		G = G_temp.read().val
		col_mean = np.nanmean(G, axis=0)
		indices = np.where(np.isnan(G))
		G[indices] = np.take(col_mean, indices[1])
		# G_S = (G - 2*f) / (np.sqrt(2 * f * (1-f)))
		#del G
		effects = np.round(np.random.normal(0,1,len(variants)),6)
		g = G@effects #g = G_S@effects
		e = np.random.normal(0,np.std(g) * np.sqrt((1-her**2)/her**2),1_000)
		y = g + e
		df_pheno = pd.read_csv(fam_path, index_col=None, header=None, delim_whitespace=True)
		df_pheno = df_pheno.rename(columns={0: "FID", 1: "IID", 2: "Father", 3: "Mother"})
		df_pheno = df_pheno[['FID','IID','Father','Mother']]
		df_pheno['g'] = np.round(g,6)
		df_pheno['e'] = np.round(e,6)
		df_pheno['y'] = np.round(y,6)
		df_pheno = df_pheno.set_index('FID')
		df_pheno.to_csv(out_pheno, sep=" ")

		df_bim = pd.read_csv(bim_path, index_col=None, header=None, delim_whitespace=True)
		df_bim_filtered = df_bim[df_bim[1].isin(variants)]

		df_causals_initial = pd.DataFrame(data={'ID':df_bim_filtered[1], 'Ref': df_bim_filtered[4], 'Beta':effects})
		df_causals_initial = df_causals_initial.set_index('ID')
		df_causals_initial.to_csv(out_causal % 0, sep=" ")
		del df_bim_filtered, df_causals_initial

		df_index = pd.read_csv(index_path, header=0, delim_whitespace=True)
		causal_index_list = random.sample(df_index['Index'].tolist(), rep)
		del df_index

		df_prs_initial = df_pheno[['IID','Father','Mother', 'g']]
		df_prs_initial.to_csv(out_prs % 0, sep=" ")
		del df_prs_initial

		del G
		G_full = snpreader.read().val
		col_mean_full = np.nanmean(G_full, axis=0)
		indices_full = np.where(np.isnan(G_full))
		G_full[indices_full] = np.take(col_mean_full, indices_full[1])

		for i_causal,j in zip(causal_index_list,range(1,len(causal_index_list)+1)):
			G_1 = G_full[:,i_causal*1_000 : (i_causal+1)*1_000]
			effects_1 = np.linalg.inv(G_1)@g # effects_1 = np.linalg.inv(G_1)@G_S@effects
			df_causals = pd.DataFrame(data={'ID':df_bim[i_causal*1_000 : (i_causal+1)*1_000][1], 'Ref': df_bim[i_causal*1_000 : (i_causal+1)*1_000][4], 'Beta':np.round(effects_1,6)})
			df_causals = df_causals.set_index('ID')
			df_causals.to_csv(out_causal % j, sep=" ")

			df_prs = df_pheno[['IID','Father','Mother']]
			df_prs['g'] = np.round(G_1@effects_1,6)
			df_prs.to_csv(out_prs % j, sep=" ")
		del df_causals, df_prs
		print('Done.')


	if args.paradox:
		print('\nSimulating a continuous trait...')

		assert args.initial_causal is not None, "Causal Markers selected by user are required for simulating a phenotype.."
		assert int(args.replicate)<5, "For datasets other than Hapmap3, we do not recommend using a replicate index greater than 5 due to computational burden. Please use replicate index < 5. Example: --replicate 4"
		
		her = float(args.h_sqrt)		
		rep = int(args.replicate)

		out_causal = args.out + ".causal"
		out_pheno = args.out + ".pheno"
		out_causal_hacked = args.out + ".causal.v%s"
		out_prs_hacked = args.out + ".prs.v%s"

		df_causals_initial = pd.read_csv(args.initial_causal, index_col=None, header=None, delim_whitespace=True)   # No column and Second column is the variant info
		causal_markers = df_causals_initial[1].tolist()

		df_variants_subset = pd.read_csv(args.geno + '.bim', index_col=None, header=None, delim_whitespace=True)
		variants_full = df_variants_subset[1].tolist()        

		assert set(causal_markers).issubset(variants_full), "selected variants to be causal markers have to be subset of variant_list.."

		snpreader = Bed(args.geno + '.bed', count_A1=True)
		

		G_full = snpreader.read().standardize().val
		
		effects_initial = np.random.normal(0,1,len(causal_markers))

		df_pheno = pd.read_csv(args.geno + '.fam', index_col=None, header=None, delim_whitespace=True)
		df_pheno = df_pheno.rename(columns={0: "FID", 1: "IID", 2: "Father", 3: "Mother"})
		df_pheno = df_pheno[['FID','IID','Father','Mother']]

		g = G_full[:,snpreader.sid_to_index(causal_markers)]@effects_initial
		e = np.random.normal(0,np.std(g) * np.sqrt((1-her**2)/her**2),G_full.shape[0])
		y = g + e

		df_pheno['g'] = g
		df_pheno['e'] = e
		df_pheno['y'] = y
		
		df_pheno = df_pheno.set_index('FID')
		df_pheno.to_csv(out_pheno, sep=" ")

		df_causals = pd.DataFrame(data={'ID':causal_markers, 'Beta':effects_initial})
		df_causals = df_causals.set_index('ID')

		df_bim = pd.read_csv(args.geno + '.bim', index_col=None, header=None, delim_whitespace=True)
		df_bim = df_bim.rename(columns={1:'ID', 4:'Ref'})
		df_bim = df_bim.set_index('ID')

		df_causals_final = pd.concat([df_causals, df_bim], join='inner', axis=1)        
		df_causals_final = df_causals_final[['Ref', 'Beta']]
		df_causals_final.to_csv(out_causal, sep=" ")
		print('Causals are uploaded..')

		del df_causals_final, df_causals, causal_markers

		assert G_full.shape[1]>1000, 'Genotype Matrix has less than 1000 columns.. Please use --hack flag in order to create paradoxical phenotype..'

		seeds = np.random.choice(list(range(int(G_full.shape[1]/20))), rep, replace=False)  # randomly choose starting seed for the first causal marker
		for seed,j in zip(seeds, range(2,len(seeds)+2)):
			index_causals = [seed]
			S = G_full[:,seed:seed+1]
			for i in range(seed+1,G_full.shape[1]+1):
				if np.linalg.matrix_rank(np.hstack((S,G_full[:,i:i+1]))) == np.linalg.matrix_rank(S)+1:
					S = np.hstack((S,G_full[:,i:i+1]))
					index_causals.append(i)

			causal_markers = [snpreader.sid[x] for x in index_causals]
			effects = np.linalg.inv(S.T@S)@S.T@g

		df_causals = pd.DataFrame(data={'ID':causal_markers, 'Beta':effects})
		df_causals = df_causals.set_index('ID')

		df_bim = pd.read_csv(args.geno + '.bim', index_col=None, header=None, delim_whitespace=True)
		df_bim = df_bim.rename(columns={1:'ID', 4:'Ref'})
		df_bim = df_bim.set_index('ID')

		df_causals_final = pd.concat([df_causals, df_bim], join='inner', axis=1)        
		df_causals_final = df_causals_final[['Ref', 'Beta']]
		df_causals_final.to_csv(out_causal_hacked % j, sep=" ")				

		df_prs = df_pheno[['IID','Father','Mother']]
		df_prs['g'] = S@effects 
		df_prs.to_csv(out_prs_hacked % j, sep=" ")


	with open(args.out + ".log", "a") as f:
		f.write("\nAnalysis End Time: %s" % datetime.datetime.now().strftime("%A - %d %B %Y - %H:%M:%S"))
		f.write("\nProgram halted in %s seconds...\n" % (round(time.time() - start_time,2)))
```
